# Quantitative genetics of trauma induced mortality in *Drosophila melanogaster*

**DOI:** 10.1101/2025.10.09.681457

**Authors:** Gwanwoou Yun, Ronchen Liu, Nathaniel P. Sharp

## Abstract

Traumatic brain injury is a major cause of chronic neurological impairment worldwide, and there is evidence that both genetic and environmental variation contribute to the likelihood of recovery. Using an insect model of traumatic brain injury, we examined variation in the risk of mortality using quantitative genetic approaches applied previously for life history traits in *Drosophila melanogaster*. We quantified additive genetic variance for mortality risk using a controlled breeding design and found levels of variation consistent with existing data on major fitness components. We found no evidence for inbreeding effects on mortality risk, suggesting that dominance genetic variance makes little contribution to this trait. To explain the high level of standing genetic variation, we considered whether mortality risk depends on the metabolic resources available to an individual, also known as “condition”. We manipulated condition by inducing random mutations and by restricting calories during larval development. We found that reduced condition due to both random mutations and resource limitation significantly increased the risk of mortality following trauma. Among inbred lines, greater mortality risk was associated with lower viability, fecundity and longevity, consistent with an effect of genome-wide genetic quality. Our results suggest that further consideration of individual condition would be valuable for understanding and predicting variation in the outcomes of traumatic brain injury.

## Introduction

Traumatic brain injury (TBI) is a significant global cause of neurological disability and mortality (Roozenbeek *et al*., 2013; Dewan *et al*., 2019). There is evidence that the outcome of TBI depends on genetic variation among individuals, with a heritability of 26% (Kals *et al*., 2022). The identification of specific loci contributing to TBI outcomes would be valuable, but genome-wide association studies have revealed that this phenotype is complex and likely influenced by variation at many loci (Gomez *et al*., 2021; Kals *et al*., 2022; Merritt *et al*., 2024). *Drosophila melanogaster* has emerged as a model system for studying TBI, allowing for high-throughput experiments investigating the genetics of this disorder (Fesharaki-Zadeh and Datta, 2024). Fly models have revealed modifiers of TBI outcomes including genetic background (Katzenberger *et al*., 2013) and age (Katzenberger *et al*., 2013), and physiological consequences of TBI such as tissue barrier dysfunction (Katzenberger *et al*., 2015), innate immunity (Katzenberger *et al*., 2013; Swanson *et al*., 2020), and sleep (Barekat *et al*., 2016; Alphen *et al*., 2022). There is a need for further study of the origin and maintenance of genetic variation that influences TBI outcomes.

Our objective was to consider TBI-induced mortality in *D. melanogaster* from a quantitative genetic perspective, using the “high-impact trauma” (HIT) model developed by Katzenberger et al. (2013). Specifically, we sought to quantify additive genetic variance for mortality rate following TBI, and to test the influence of recessive genetic variation, spontaneous mutations, and individual condition. If mortality rate is influenced by recessive alleles, we would expect the risk to differ between inbred and outbred flies. In a previous study, crosses between inbred lines with high and low mortality resulted in strains with intermediate mortality (Katzenberger *et al*., 2015), suggesting that the differences between the parental lines were driven mainly by additive genetic variation. We tested for inbreeding depression in HIT mortality by sampling from an outcrossing population, thereby incorporating a wider array of standing genetic variation.The additive genetic variation for a polygenic trait will largely determine the response to selection (Falconer and Mackay, 1996; Walsh and Lynch, 2018). Previous studies have demonstrated significant variation in HIT mortality among inbred lines (Katzenberger *et al*., 2013, 2015, 2023), which would reflect overall genetic variation (additive and non-additive). In addition to testing for dominance variance in the form of inbreeding depression as described above, we sought to quantify additive genetic variance for HIT mortality using a half-sibling breeding design (Lynch and Walsh, 1998). This approach allowed us to quantify the narrow-sense heritability for the trait and compare standardized variance values among traits. We were particularly interested in whether the level of genetic variation for HIT mortality is similar to life history traits that have been studied in flies. Additionally, we examined the HIT mortality values that have been reported previously for lines from the Drosophila Genetic Reference Panel (DGRP; Mackay *et al*., 2012), in relation to other traits that have been measured for the same lines, allowing us to assess correlations among traits.

As with any other quantitative trait, standing heritable variation in HIT mortality could be attributable, at least in part, to deleterious mutation-selection balance (Houle *et al*., 1994, 1996; Charlesworth and Hughes, 1999; Zhang and Hill, 2005; Charlesworth, 2015), with directional selection favoring lower mortality. If many loci contribute to this trait, random mutagenesis would cause increased susceptibility to mortality.Alternatively, if this trait is influenced by relatively few loci, we would be less likely to detect a significant effect of mutagenesis. If the trait is actually under stabilizing selection favoring intermediate values (e.g., due to a strong trade-off with another fitness component), selection would favor reduced variance; we would then expect random mutations to increase the trait variance, with little effect on the trait mean (Keightley and Hill, 1988). We tested the effect of random mutations by exposing male flies to a chemical mutagen and then measuring HIT mortality in their offspring.

One way to account for the high levels of genetic variance often observed in major fitness components like viability and fecundity (Houle, 1992) is to posit that these traits reflect the ability of an individual to acquire resources, which in turn is depends on many loci (Houle, 1991; Rowe and Houle, 1996; Tomkins *et al*., 2004; Hooper and Bonduriansky, 2022). The “condition” of an individual can be affected by its genetic quality, but also by the availability of resources in its environment, though these influences may not be entirely concordant (Clark *et al*., 2012; Bonduriansky *et al*., 2015; Hooper and Bonduriansky, 2022). We tested for an effect of environmentally induced variation in condition by rearing flies in high-or low-quality larval environments and measuring their susceptibility to HIT-induced mortality as adults.We find that HIT mortality shows high levels of additive genetic variation and condition-dependence. Correlations with other traits suggest that HIT mortality is influenced by many loci that also contribute to variation in other fitness components, such that variation in this trait is maintained by a balance between mutation and purifying selection.

### Materials and Methods

#### Fly strains and culturing

We conducted all experiments with flies from the Canton-S wild-type genetic background, obtained from the Bloomington Drosophila Stock Center (RRID:BDSC_64349) and maintained as a large outbreeding cage population with overlapping generations since 2019. We maintained flies at 25 C, under a 12:12 light:dark cycle, and conducted crosses under CO_2_ anesthesia using flies that were 2-5 days old post eclosion, with females collected as virgins. Except where noted, we cultured flies on defined medium (14.3 g/L agar, 92.3 g/L white sugar, 46 g/L debittered yeast, 7.4 g/L potassium sodium tartrate, 0.93 g/L potassium phosphate, 0.46 g/L sodium chloride, 0.46 g/L calcium chloride, 0.46 g/L magnesium chloride, 0.46 g/L iron sulfate, 0.5% propionic acid) in standard vials seeded with live yeast.

#### High-impact trauma and mortality index

We subjected flies to high-impact trauma (HIT) following an approach described in detail previously (Katzenberger *et al*., 2013). For each replicate, we transferred a group of flies to an empty vial without anesthesia and restricted them to the bottom quarter of the vial with a cotton ball. We then attached the vial to the free end of a metal spring clamped to a wooden board, deflected the spring at a right angle and released it, causing the vial to contact a polyurethane pad at approximately 3 m/s (Katzenberger *et al*., 2013). For each replicate vial we applied this procedure three times, separated by 5 min of recovery time. We then transferred flies into vials with media, incubated them for 24 h, and then scored the number of dead flies in each vial. In addition to vials of flies subjected to trauma, we generated control vials where matched flies were transferred into empty vials for the same amount of time as the treatment flies but were not subjected to trauma. This “sham” treatment allowed us to control for any variation in mortality not caused by trauma. For each experimental unit (e.g., flies from the same family), we calculated the mortality index (MI) as the proportion of dead flies in the treatment vial minus the proportion of dead flies in the sham vial, multiplied by 100. Our experiments vary somewhat in the number of flies per treated vial, and combine males and females, but there is evidence that these factors do not influence mortality (Katzenberger *et al*., 2013).

#### Genetic variance

We used a half-sibling breeding design to estimate genetic variance in mortality following high-impact trauma, using flies collected at random from the Canton-S population. We first created multiple “families” with one male (“sire”) and three virgin females (“dams”) each. Following one day of mating, we discarded the males and placed the females in individual oviposition vials. From each oviposition vial we collected adult offspring and subjected half to the HIT treatment and half to the sham treatment, with an average of 79 flies per MI measure. Some females did not produce offspring; excluding such cases, this experiment was ultimately comprised of 55 sires, 133 dams, and 10496 offspring, tested in two roughly equal blocks. We calculated MI for the offspring of each dam and determined the among-sire component of variance by fitting a linear mixed model using the *R* package lme4 (Bates *et al*., 2015). In a half-sibling breeding design, the additive genetic variance is given by four times the among-sire component of variance, assuming epistasis is negligible (Lynch and Walsh, 1998).

#### Inbreeding depression

To test for an effect of inbreeding on mortality following high-impact trauma, we first collected flies at random from the Canton-S population and created vials with one male and one virgin female each. From each of these “families” we collected male and female offspring as virgins. We then generated outbred flies by crossing males from one family with females from a different family; we generated inbred flies by crossing males and females from the same family (full siblings). We were ultimately able to obtain MI data for 34 outbred and 34 inbred groups of flies, with an average of 49 flies per MI measure (3331 flies in total). We tested for an effect of inbreeding on mean MI using a t-test and compared coefficients of variation using a randomization test with 10000 replicates.

#### Mutagenesis

To test the impact of random mutations on mortality following high-impact trauma, we first collected random males and virgin females from the Canton-S population. We placed groups of males in empty vials for two hours to induce starvation and then transferred them into bottles containing filter paper saturated with either sugar water (1 g/mL sucrose), or sugar water with 3 mM methyl methanesulfonate (MMS). MMS is an alkylating agent and mutagen, which is known to generate point mutations and aberrations in flies (Ashburner *et al*., 2005). After 24 h, we combined treated males with virgin females, with one male per vial. We expect offspring from these vials to be wild type when sired by control males, or to be heterozygous for induced mutations when sired by mutagenized males. We were ultimately able to obtain MI data for 25 control and 25 mutagenized groups of flies, with an average of 46 flies per MI measure (2316 flies in total). We tested for an effect of mutagenesis on MI using a t-test.

#### Larval diet quality

To test the impact of larval diet quality on mortality following HIT, we first placed oviposition plates in the Canton-S cage overnight. Following egg hatching, we picked L1 larvae and placed them in groups of 50 in vials of our standard media (described above; “high quality” diet), or in vials containing media with half of the standard amounts of sugar and yeast (“low quality” diet). We collected adults emerging in these vials and preserved some for body mass measurements (*N* = 51 flies per sex and diet treatment) and used others for MI measurements. To measure body mass, we dried flies at 70 C for 24 h and weighed them individually to the nearest 0.01 mg. We tested for effects of larval diet on body mass using t-tests (Welch t-test in the case of males, where Levene’s test indicated unequal variances).

In this assay, flies were collected in mass across vials in each treatment, and so there was no pairing of HIT vials and sham vials. We ultimately obtained mortality data for 38 hit vials and 30 sham vials from the high-quality diet treatment, and for 44 HIT vials and 43 sham vials from the low-quality diet treatment, with an average of 29.8 flies per vial (4624 flies in total). We calculated MI for each HIT vial by subtracting the average mortality across all sham vials from the corresponding diet treatment. We tested for an effect of diet quality on MI using a t-test. An alternative analysis of mortality in all HIT and sham replicates using a quasi-binomial generalized linear model produced the same conclusions.

#### Trait correlations

The rate of mortality following HIT has previously been investigated for inbred strains from the DGRP (Katzenberger *et al*., 2015), where a variety of other traits have also been measured. To understand the relationship between mortality from trauma and underlying genetic quality, we obtained publicly available data on additional life history traits from these same lines. Specifically, we considered data on survival to adulthood (viability), using the average across three temperatures tested (Ellis *et al*., 2014), and data on lifespan and lifetime fecundity, using values corrected for body size and block (Durham *et al*., 2014). Of the 179 DGRP lines with MI data, data on viability were available for 49 cases, and data on lifespan and fecundity were available for 169 cases. We calculated Pearson correlation coefficients for pairs of traits and tested for significance using bootstrapping with 10000 replicates.

## Results

### High levels of genetic variance

We detected significant additive genetic variance in HIT mortality in the Canton-S laboratory population (χ^2^ = 10.31, *df* = 1, *P* = 0.001; Table S1). We estimate a coefficient of additive genetic variation (Houle, 1998) of 46.2%, and a narrow sense heritability of68.4%. The range of mean MI by sire we observed (Fig. 1) is consistent with the range observed for DGRP lines (Katzenberger *et al*., 2015). For comparison, we calculated the coefficient of total genetic variance for HIT mortality among the DGRP inbred lines studied by Katzenberger *et al*. (2015) and found a value of 36.9%. Thus, two studies of different fly populations using different methods both indicate relatively high levels of standing genetic variation for HIT mortality.

**Figure 1.**
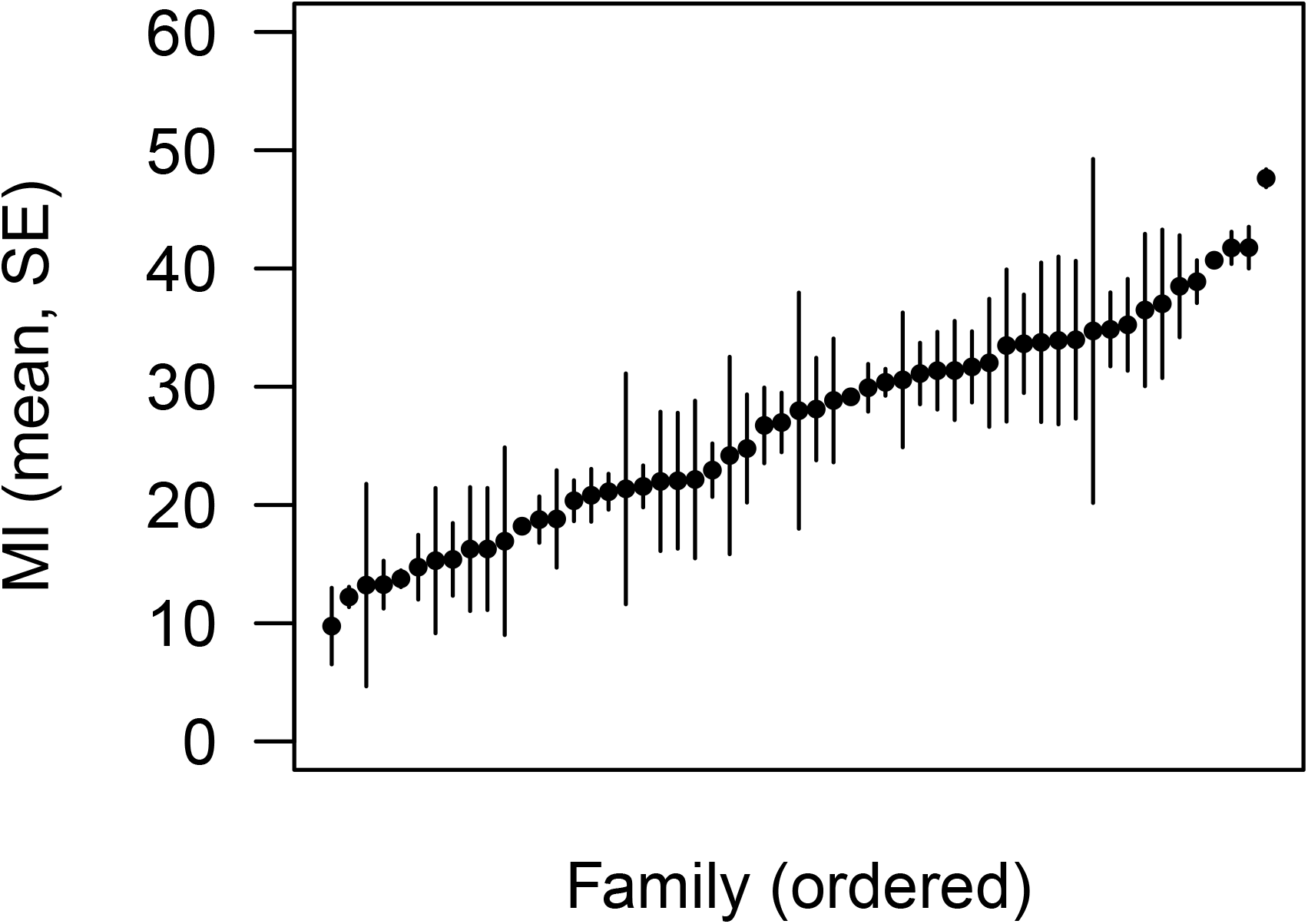
Genetic variance in mortality following high-impact trauma. Points represent means for 55 families of the mortality index (MI), ordered from low to high. We detected significant additive genetic variance in MI.

### No detectable inbreeding depression

Inbreeding could affect mortality following TBI if there are recessive alleles contributing to variation in this trait, i.e., dominance genetic variance (Charlesworth and Willis, 2009). Our analysis comparing MI in families derived from full-sibling mating (inbred) and outbred families (Fig. 2; Table S2) revealed no effect of inbreeding on the trait mean (*t* = 0.69, *df* = 66, *P* = 0.49), although there was significant HIT-induced mortality overall (*t* = 6.17, *df* = 67, *P* = 4.56 × 10^-8^). The coefficient of variation in MI was greater in inbred families than in outbred families (158.4% versus 114.7%), but this difference was not statistically significant (randomization, *P* = 0.31). The observation that inbreeding did not significantly affect the mean or variance in HIT mortality suggests that most genetic variants affecting HIT mortality do so in an additive manner.

**Figure 2.**
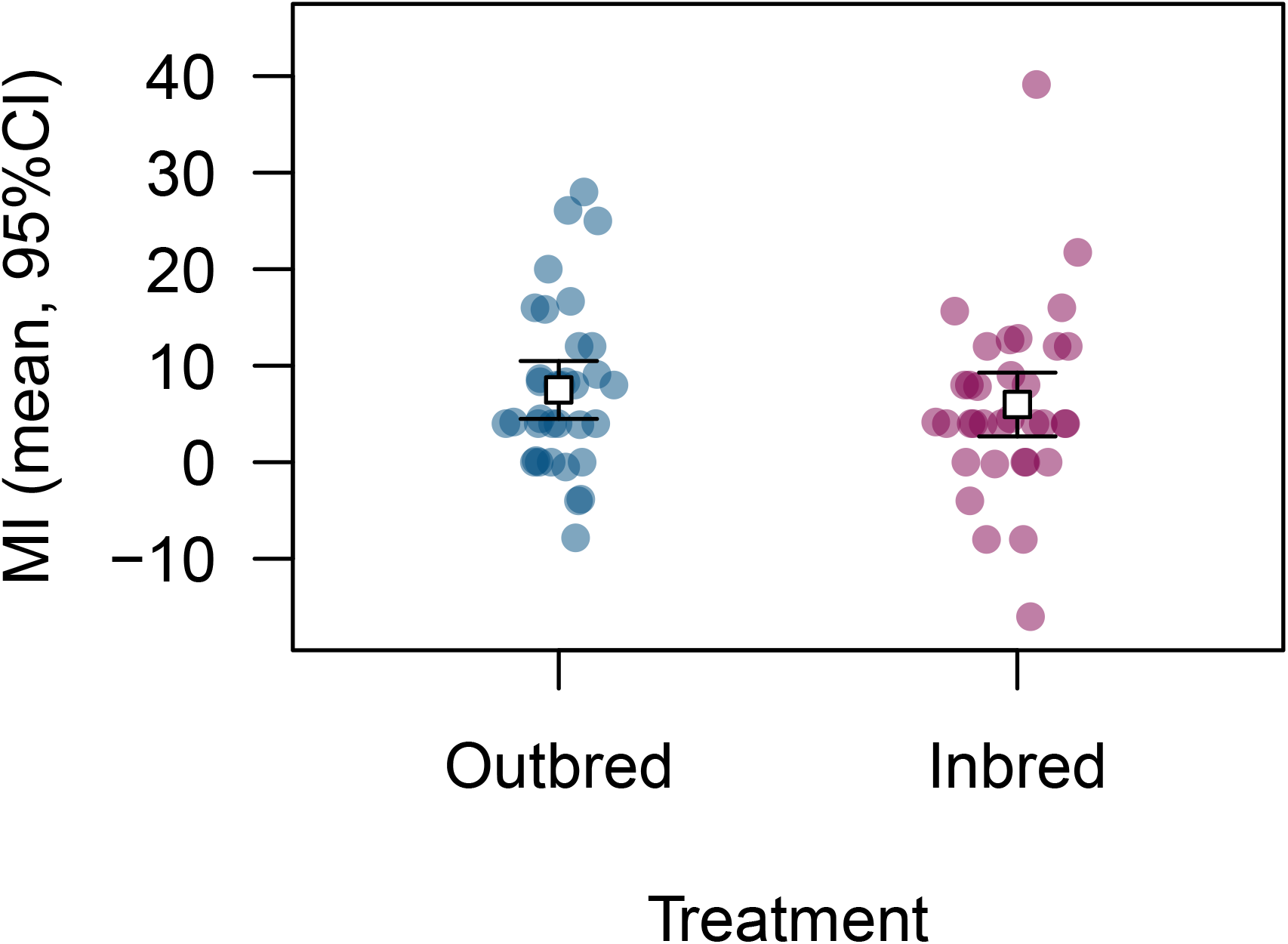
Effect of inbreeding on mortality following high-impact trauma. Inbred flies were the result of mating between full siblings; outbred families were the result of mating between unrelated individuals. We did not detect a significant effect of inbreeding on MI.

### Mutagenesis increases HIT mortality

Standing genetic variation in mortality following HIT could be maintained in part by mutation-selection balance if this trait is a large “mutational target”. We found that the average MI was significantly higher (8.8%) in mutagenized families than in non-mutagenized families (Fig. 3; Table S3; *t* = –2.39, *df* = 48, *P* = 0.021). We did not detect an effect of mutagenesis on the coefficient of variation for MI (randomization with 10000 replicates, *P* = 0.19), though existing genetic variation could have partially obscured a signal of increased variation following mutagenesis. The dose of MMS we used, which has been found to have a relatively weak effect on heterozygous viability (Melde *et al*., 2024), was nevertheless sufficient to significantly affect HIT mortality, suggesting that many loci contribute to the outcome of TBI.

**Figure 3.**
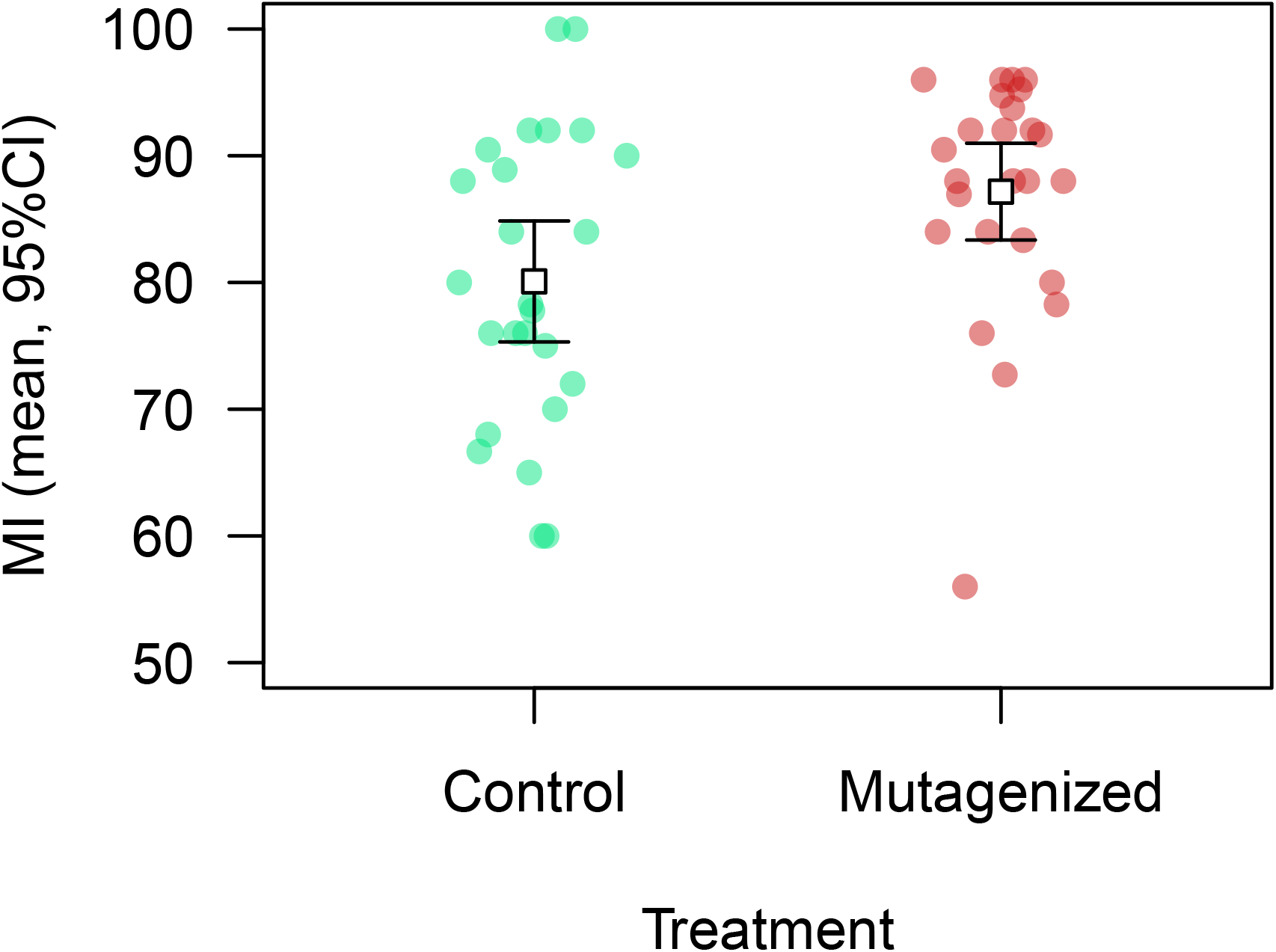
Effect of mutagenesis on mortality following high-impact trauma. Mutagenized families were derived from males exposed to starvation followed by sugar water containing the mutagen MMS; control families were derived from males exposed to starvation followed by sugar water only. Mutagenesis resulted in a significant increase in MI.

### Low quality larval diet increases HIT mortality

Just as random mutations could influence HIT mortality by affecting the ability of individuals to acquire resources, development in a resource-limited environment could also influence the outcomes of trauma. We reared flies on either standard media or media with half of the usual amount of yeast and sugar; we found that the low-quality diet reduced female body mass by 20% (Table S4; *t* = 8.09, *df* = 100, *P* = 1.48 × 10^-12^) and reduced male body mass by 16% (Table S4; Welch *t* = 8.25, *df* = 83.23, *P* = 2.00 × 10^-12^). By itself, we would expect a reduction in body mass to lead to a lower rate of HIT mortality, since smaller flies experience less force during the HIT procedure (Johnson-Schlitz *et al*., 2022). However, we found that average MI was 3.5-fold higher in flies reared on the low-quality larval diet (Fig. 4; Table S5; *t* = 14.9, *df* = 80, *P* < 2.2 × 10^-16^), indicating that HIT mortality is highly sensitive to variation in resource availability during larval development.

**Figure 4.**
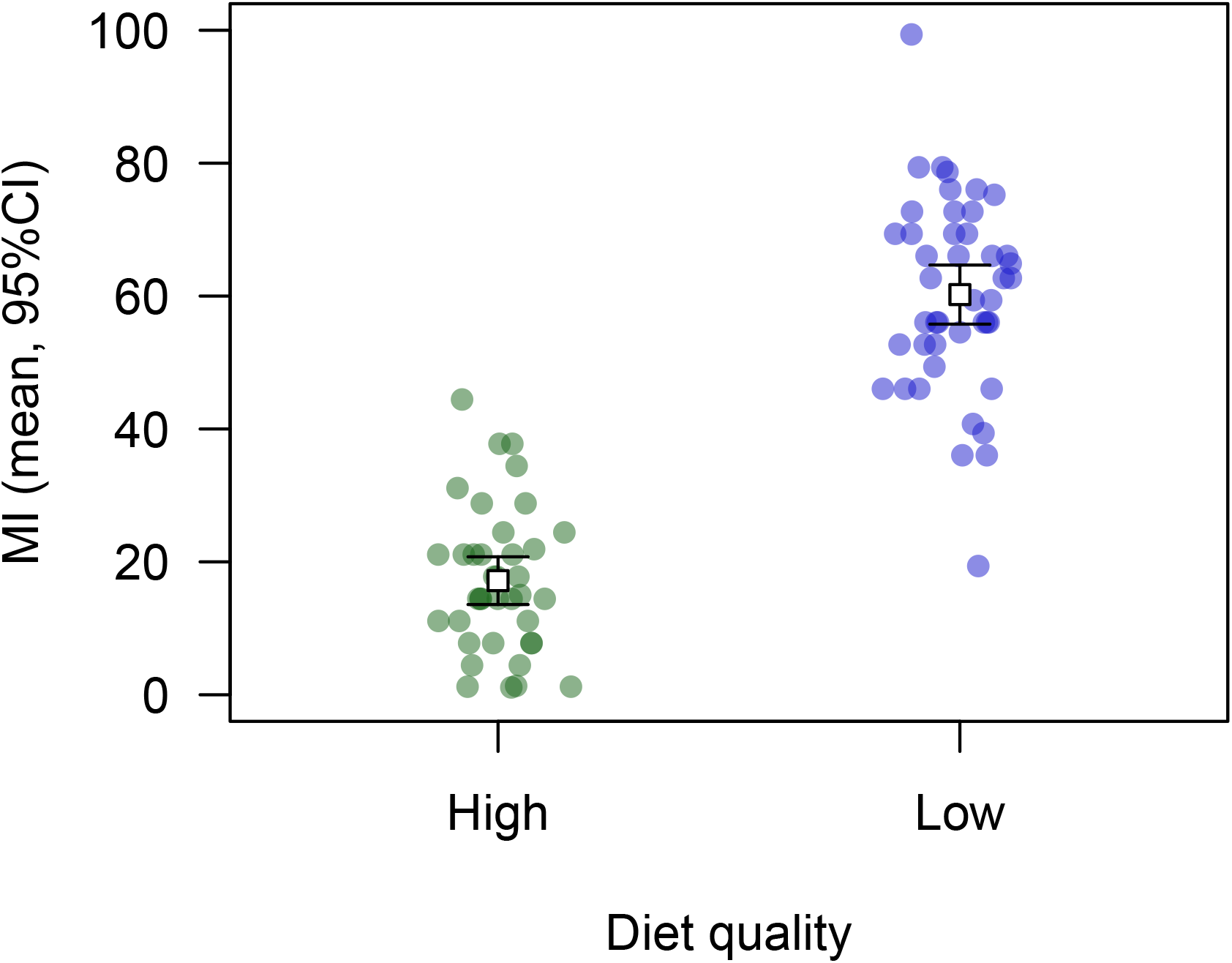
Effect of larval diet quality on mortality following high-impact trauma. Flies reared from the L1 larval stage on a low-quality diet (half of the standard amount of sugar and yeast) displayed higher MI than flies reared on a standard diet.

### HIT mortality is correlated with fitness components

Our findings above suggest that mortality following HIT is affected by random mutations and condition, which are underlying variables that are also expected to affect fitness related traits. We therefore predicted that HIT mortality would be negatively correlated with fitness-related traits across genotypes. Based on analyses of sets of DGRP lines where both MI and fitness information is available, we confirmed this prediction (Fig. 5). Specifically, we found that MI was negatively correlated with egg-to-adult viability (*r* = – 0.28, bootstrap *P* = 0.029), lifetime fecundity (*r* = –0.20, bootstrap *P* = 0.010), and lifespan (*r* = –0.20, bootstrap *P* = 0.002). The mortality risk in HIT-treated flies is therefore greater in genotypes that have lower fitness in the absence of HIT.

**Figure 5.**
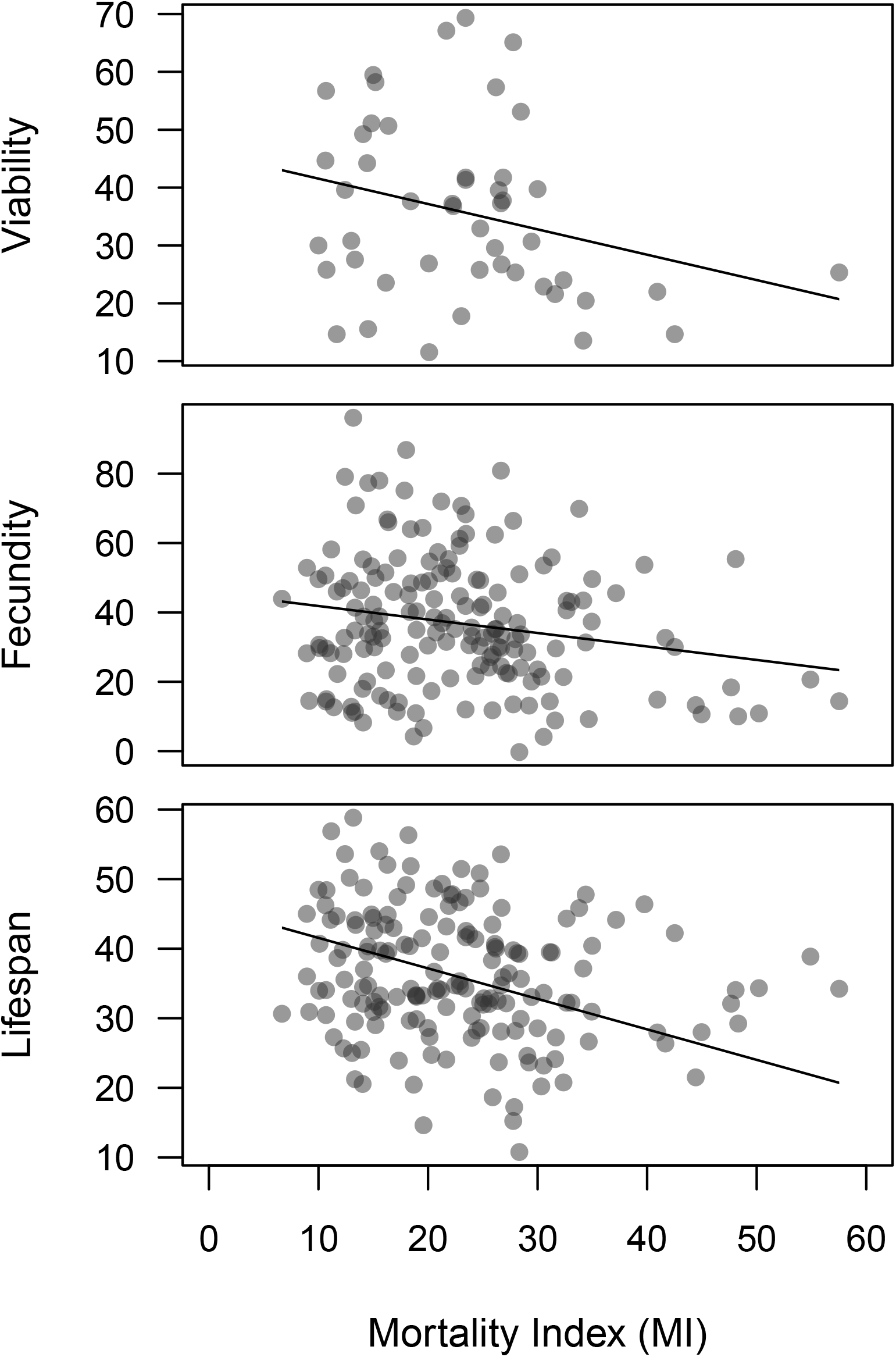
Relationships between fitness-related traits and mortality following high-impact trauma. Analysis of published data on DGRP lines (Durham *et al*., 2014; Ellis *et al*., 2014; Katzenberger *et al*., 2015) revealed significant negative correlations between MI and viability (top), fecundity (middle) and lifespan (bottom). Linear model trendlines are shown in each panel.

Previous investigations of the relationship between MI and genotype in the DGRP lines revealed a common genetic variant in the gene *grh* which was associated with elevated mortality following HIT (Katzenberger *et al*., 2015). To determine whether the above trait relationships persist in the absence of this variant, we repeated our analyses excluding DGRP lines harboring *grh* variants and obtained similar results for viability (*r* = –0.28, bootstrap *P* = 0.056), fecundity (*r* = –0.21, bootstrap *P* = 0.008) and lifespan (*r* = –0.24,bootstrap *P* = 2 × 10^-4^). Variation at the *grh* gene therefore does not account for the overall genetic correlation between HIT mortality and fitness-related traits.

## Discussion

We used an established insect model to investigate the quantitative genetics of mortality following traumatic injury. There appears to be substantial additive genetic variation for this trait (Fig. 1), but we did not find evidence for inbreeding depression (Fig. 2); considering the coefficient of additive genetic variation, which uses the trait mean for standardization, genetic variation in HIT mortality (46.2%) is within the range of previous reports on *Drosophila* life history traits like male mating success, female fecundity, and larval viability (Houle, 1992, 1998; Charlesworth and Hughes, 1999; Sharp and Agrawal, 2018). This suggests that variation in HIT mortality has a broad genetic basis, as with major fitness components.

Our inbreeding test for dominance variance would not be effective if alleles affecting HIT mortality are disproportionately likely to be recessive lethal, since such alleles would be purged prior to our testing. Previous studies indicate that recessive lethality cannot be the sole cause of inbreeding depression for life history traits in *Drosophila* (Latter *et al*., 1995; Fry *et al*., 1998; Charlesworth *et al*., 2007; Mallet and Chippindale, 2011). It is unclear whether HIT mortality is an exception to this pattern. A more likely explanation is that we lacked the statistical power to detect inbreeding depression in this case; simulations using the variance in MI observed in this part of our study indicate that we would have <50% power to detect a true increase in MI due to inbreeding of 60% or less. We also observed lower overall mortality here (Fig. 2) than in other parts of our study, though it was still significant. We therefore cannot rule out the presence of mild or moderate inbreeding depression for HIT mortality, which might be detectable using larger samples or higher levels of trauma.

Laboratory populations may not always be representative of natural populations, and so we compared our genetic variance estimate with data from Katzenberger *et al*. (2015), who characterized variance among inbred DGRP lines. If there is little dominance variance for this trait in the DGRP population (as appears to be the case in our lab population), then the variance among DGRP lines should mainly reflect additive genetic variation (or additive-by-additive epistasis). We calculated a standardized genetic variance among DGRP lines is 36.9%, similar to what we observed. Lower genetic variance in the DGRP lines would be expected if HIT mortality is affected by recessive deleterious alleles that were purged during inbreeding; crosses between DGRP lines with very different trait values resulted in progeny with trait values near the midparent value (Katzenberger *et al*., 2015), suggesting a lack of dominance effects following inbreeding. We conclude that there is a relatively high level of genetic variation for mortality following traumatic injury in flies, in both laboratory and wild populations, and that much of this variance is likely additive.If HIT mortality acts as a condition-dependent trait, the risk of mortality could increase following random mutagenesis or diet restriction. We found that flies that inherited mutagenized chromosomes were significantly more susceptible to mortality following HIT (Fig. 3), suggesting that this trait is a large mutational target (Houle *et al*., 1996; Houle, 1998), and that natural selection has historically favored alleles whose effects include reduced HIT mortality. In our study, the flies tested would carry induced mutations in the heterozygous state, with the exception of any Y-linked mutations; although we did not detect inbreeding depression involving standing variance, some induced mutations could still have recessive effects on HIT mortality that would be revealed in homozygous flies.

We also found that flies reared on a low-quality larval diet were substantially more susceptible to mortality following HIT (Fig. 4). Our diet manipulation reduced adult dry body mass by 16-20%; for comparison, a diet manipulation causing a similar body mass reduction detectably reduced male sperm competition success (Clark *et al*., 2012). The increase in HIT mortality in the presence of both random mutations and reduced larval diet quality suggests that HIT mortality is condition dependent. This is concordant with results from rodent models, which indicate that negative TBI outcomes are exacerbated by early life stress (Fesharaki-Zadeh *et al*., 2020; Sanchez *et al*., 2021).If HIT mortality depends on genetic quality, we would predict that genotypes that are more susceptible to HIT would also show reduced values of key life history components. We were able to confirm this prediction by examining trait correlations for sets of DGRP lines: higher HIT mortality was associated with lower viability, fecundity, and lifespan (Fig. 5). There is also evidence that older flies are more susceptible to HIT (Katzenberger *et al*., 2013), consistent with the increased expression of deleterious alleles with age predicted under the mutation accumulation hypothesis for the evolution of aging (Charlesworth, 2000; Li *et al*., 2023).

Taken together, our findings indicate that flies with reduced condition––either because of deleterious genetic variants or because of reduced access to resources during development––are at increased risk of mortality following traumatic injury. In humans, there is growing interest in the complex ways that social, demographic and environmental factors may influence recovery from TBI (Ponsford, 2013; Haarbauer-Krupa *et al*., 2021). We suggest that further consideration of individual condition alongside specific risk factors may be a useful framework for explaining and predicting variation in TBI outcomes.

## Acknowledgements and funding sources

Thanks to P. Nelson for assistance in the lab. Thanks to D. Wassarman for helpful feedback and for providing the HIT device used in this study. Fly mass data were obtained at the University of Wisconsin–Madison Biophysics Instrumentation Facility, established with support from grants BIR-9512577 (NSF) and S10 RR13790 (NIH). Research reported in this publication was supported by the National Institute of General Medical Sciences of the National Institutes of Health under award number R35GM154954 to N.S.

## Author contribution statement

N.S. designed the experiments. G.Y., R.L. and N.S. performed the experiments. N.S. analyzed the data and wrote the manuscript.

## Conflict of Interest

The authors declare that no conflict of interest exists.

## Data archiving

The data associated with this study are available as Supplementary Material.

## Notes

### Competing Interest Statement

The authors have declared no competing interest.

